# ORGANOID-BASED ASSESSMENT OF ENHANCER ACTIVITY VIA RECOMBINATION-MEDIATED TRANGENESIS

**DOI:** 10.64898/2026.11.13.766570

**Authors:** Jorge Mañes-García, Diego Martin-Sanz, Eva Pajda-Szeligowska, Raquel Marco-Ferreres, Noemi Tabanera, Elvira Zakirova, Silvia K Nicolis, Paola Bovolenta, Leonardo Beccari

## Abstract

Functional characterization of cis-regulatory elements (CREs) is essential for understanding developmental gene regulation and interpreting human noncoding variation. However, conventional transgenesis assays are limited by integration copy number variability and genomic position effects, while species-specific developmental differences constrain the study of human regulatory sequences in classical model organisms, demanding assays based on human-specific contexts.

Here, we developed RepoV, a transgenesis vector enabling single-copy integration of enhancer reporter constructs in the genome of human iPS cells through PhiC31-mediated recombination. Transgenic iPSCs were subsequently used to assess candidate neurodevelopmental enhancers in neural organoids. RepoV recapitulated the expected activity of previously characterized regulatory elements and enabled the evaluation of candidate sequences associated with the neurodevelopmental genes *POU3F2* and *FOXG1*. Notably, two *FOXG1* enhancers displayed different activities in zebrafish larvae and human neural organoids. The organoid results were more consistent with *endogenous the FOXG1* expression and the pathological phenotypes associated with disease-related regulatory variants identified in patients. Together, these findings establish RepoV transgenesis combined with human organoid cultures as a valuable platform for the functional evaluation of human developmental regulatory sequences.

## INTRODUCTION

Coordination of gene expression across space and time underlies cell identity, development, and tissue homeostasis in multicellular organisms. Much of this control is mediated by cis-regulatory elements (CREs): DNA sequences, typically 200bp −2kb in length, that recruit combinations of transcription factors to activate or repress gene activity (Spitz 2016). They can be located in intergenic, intronic, or exonic regions and regulate target genes over distances exceeding 1 megabase through chromatin looping (Mouren et al. 2026; Spitz 2016). Developmental genes, in particular, are often controlled by complex regulatory landscapes in which multiple CREs act cooperatively or redundantly to generate dynamic and tissue-specific expression patterns (Andrey et al. 2013; Harmston et al. 2017; Hörnblad et al. 2021; Lettice et al. 2003). Genetic variants and structural rearrangements affecting CREs relate with a wide range of developmental disorders, metabolic diseases, and cancer (Collins et al. 2020; Doan et al. 2016; Jeong et al. 2008; Lupiáñez et al. 2015), and over 90% of disease-associated sequence variants are estimated to lie in the non-coding genome (Khurana et al., 2016; Schaub et al., 2012). Besides, changes in gene regulation have likewise contributed to metazoan evolution (Bi et al. 2023; Prescott et al. 2015; Reilly et al. 2015).

Genome-wide epigenomic approaches have identified millions of putative CREs on the basis of chromatin accessibility and epigenetic marks (ENCODE Project Consortium 2012; Gao and Qian 2020; Heintzman et al. 2007; Visel et al. 2009). A large fraction of these are enhancers (Bulger and Groudine 2010; ENCODE Project Consortium 2012): elements that promote transcription of their target genes largely independently of distance and orientation (Banerji et al. 1981). However, while epigenetic profiling predicts elements with regulatory potential, it provides limited information about CRE’s function or how sequence variants impact their activity and cognate gene expression. Consequently, functional characterization of CREs remains a major challenge of the post-genomic era.

Transgenesis assays constitute the main functional approach for assessing enhancer activity in vivo (Kvon 2015). Candidate sequences are cloned upstream of a reporter gene and the construct is integrated into the genome of the one cell-stage embryo of a chosen model organism. Classical approaches rely on random integration through the cell endogenous DNA repair pathways or on transposase-or lentivirus-mediated insertion (e.g. (Bessa et al. 2009; Fong and Ceroni 2025; Friedli et al. 2010; Zelensky et al. 2017). However, interpretation of transgene activity in these experiments is hampered by the variability in transgene copy number (affecting reporter expression levels (Dorer 1997; Garrick et al. 1998; Sabl and Henikoff 1996)) and in genomic insertion sites which can affect transgene expression via either positive or negative genome position effects (Robertson et al. 1995; Clark et al. 1994). More recent targeted integration strategies based on CRISPR or site-specific recombinases have improved reproducibility by directing single-copy insertion to defined genomic loci (Kvon et al. 2020; Osterwalder et al. 2022) but are generally limited by insert size or require engineered genomic acceptor sites, respectively (Chen et al. 2024). Together, these technical constraints complicate the interpretation of enhancer activity and limit assay scalability, ultimately hampering the systematic analysis of CRE function.

The choice of model organism represents an additional challenge. Orthologous enhancers from different species can display either similar or divergent regulatory activity despite their sequence conservation (e.g.(Gehrke et al. 2015; Guerreiro et al. 2016; Hintermann et al. 2022; Ritter et al. 2010; Villar et al. 2015; Yang et al. 2015)). This functional variability, together with the developmental and anatomical differences separating distantly related species, complicates the interpretation of enhancer activity in cross-species assays. Selecting an appropriate model therefore requires balancing its phylogenetic proximity to the species from which the candidate sequence derives against experimental and ethical considerations associated with animal research. Murine models, for instance, offer a high degree of genetic and developmental conservation with human, but transgenesis is laborious, costly, and demands large numbers of animals. Instead, other vertebrate models such as zebrafish or Xenopus, enable rapid and efficient transgenesis, but at the cost of greater phylogenetic divergence.

The human cerebral cortex has undergone extensive changes in developmental and cis-regulatory programs, even relative to closely related primates (Emera et al. 2016; Kozlenkov et al. 2020; Reilly et al. 2015) and numerous studies have linked alterations in the regulatory landscapes of neurodevelopmental genes to brain disorders and with cognitive or behavioral traits (D’haene and Vergult 2021; Doan et al. 2016; Jeong et al. 2008; Short et al. 2018). Human brain organoids have emerged as powerful experimental systems that recapitulate key aspects of human corticogenesis and have enabled important advances in the study of cortical development, evolution, and disease (reviewed in (Eichmüller and Knoblich 2022; Pollen et al. 2019).

We combined the use of human brain organoids with the amenability of human induced pluripotent stem cells (iPSCs) for genetic engineering, to establish an efficient platform for functional evaluation of human CREs. For that, we developed RepoV, a PhiC31 integrase-based enhancer transgenesis system that drives stable single-copy integration of reporter constructs into endogenous *attP* sites in human iPSCs. After selection, transgenic lines are differentiated directly into brain organoids, allowing multiple candidate enhancers to be analyzed in parallel. Combining this strategy with region-specific neural organoids, allows to compare regulatory activities across distinct neurodevelopmental contexts. As proof of principle, we validated the system using previously described enhancers linked to the human *EMX1*, *NR2F1* and of the medaka *Rx2* genes. Finally, we characterized the activity of four putative CREs of the human *FOXG1* and *POU3F2* genes, showing that this approach could constitute a powerful tool to bridge the current gap among the high number of putative regulatory elements annotated in the human genome and their functional characterization.

## RESULTS

### RepoV plasmid construction

Site-specific recombinases constitute essential tools for targeted transgenesis in mammalian cells, enabling controlled integration of transgenes into defined genomic loci (Tasic et al. 2011). The most widely used systems include the Cre and FLP tyrosine recombinases, as well as large serine recombinases such as the PhiC31 integrase. Serine integrases mediate largely irreversible recombination between distinct attB and attP sites, making them particularly well suited for stable, single-copy transgene insertion. PhiC31 integrase in particular has been used for efficient transgenesis in different species (Chalberg et al. 2006a; Mosimann et al. 2013; Sekhavati et al. 2018) because it can target endogenous pseudo-attP sites of the genome (Chalberg et al. 2006b), removing the need to first engineer attP landing sites.

To develop a PhiC31-mediated transgenesis vector that minimizes position effects from the genomic environment at integration sites, we drew on the Zebrafish Enhancer Detection (ZED) plasmid (Bessa et al. 2009). ZED carries a dual-reporter cassette in which the candidate CRE is cloned upstream of GFP, while a tdTomato reporter driven by the cardiac actin promoter acts as a positive control for transgenesis in zebrafish larvae. Critically, the GFP cassette is flanked by the well-characterized HS4 and GAB insulators, that shield it from effects of the surrounding genome.

Our RepoV vector (**Fig. 1A**) follows the same design principle, and carries a nuclear enhanced yellow fluorescent protein (nuEYFP) reporter driven by the β-globin minimal promoter (bMP) and flanked by HS4 and GAB insulators. To minimize vector size, a single neomycin phosphotransferase II (neo/NPTII) cassette is placed under both the eukaryotic SV40 and bacterial ampicillin promoters, conferring resistance to G418 in mammalian cells and to kanamycin in bacteria. A single PhiC31 attB site enables site-specific integration into endogenous genomic attP sites (**Fig. 1A**). After integration, transgenic iPSC clones can be selected by G418 resistance (NeoR), isolated, expanded, and used to generate organoids for enhancer activity assessment (**Fig. 1B**). Alternatively, polyclonal transgenic populations can be used directly for organoid generation, for a faster readout. Candidate CREs can be cloned upstream of the bMP by TA cloning following vector linearization with XcmI digestion, which excises a ccdb cassette that serves as negative selection for undigested or partially digested vector in bacteria. Alternatively, candidates can be cloned by Gibson assembly (Gibson et al. 2009), providing a flexible platform for inserting candidate CREs or combinations thereof into RepoV.

**Figure 1.**
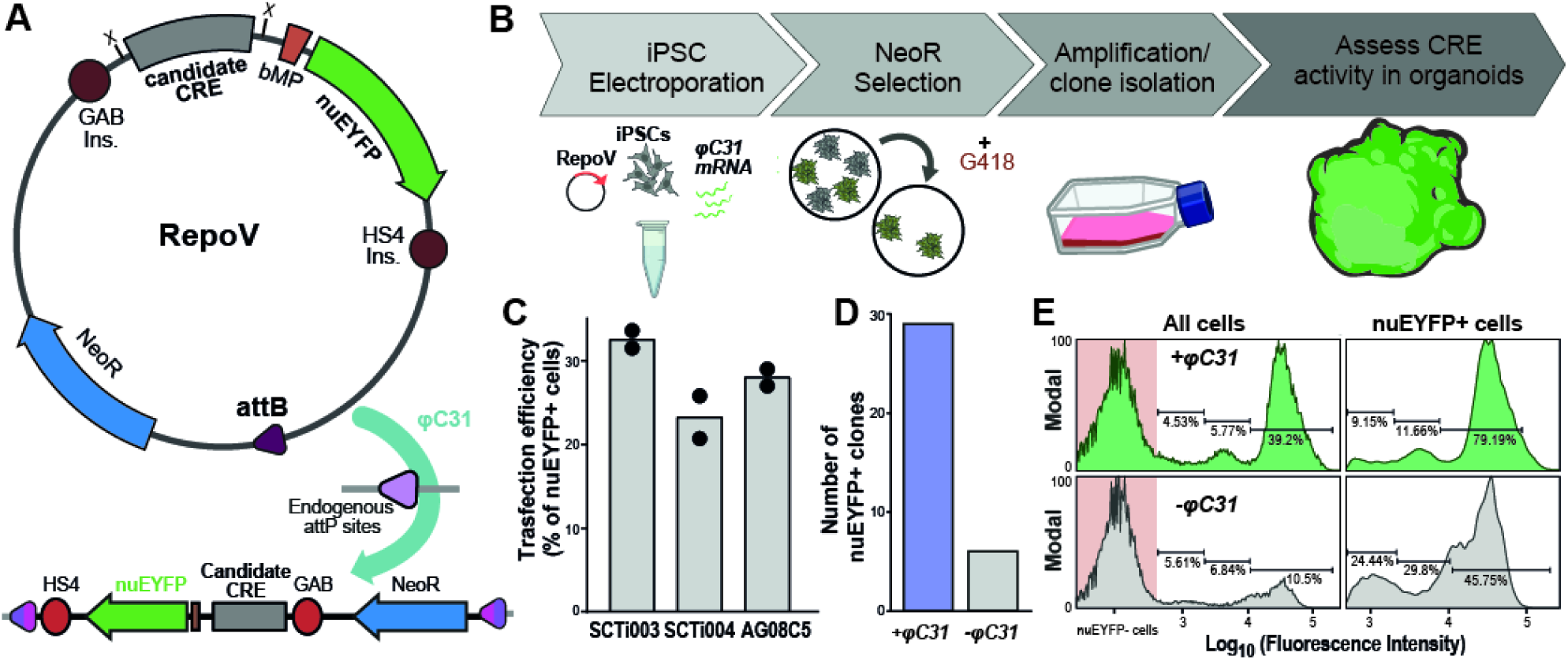
PhiC31-mediated transgenesis in human iPSCs. **A.** Scheme of the RepoV plasmid and its PhiC31-mediated integration into endogenous attP sites of the human genome. **B.** Experimental procedure used for establishing transgenic iPSC lines and evaluate enhancer activity in organoid cultures. **C.** Bar graph showing electroporation efficiency in different iPSC lines. **D.** Bar graph quantifying the number of stable transgenic clones obtained electroporating iPSCs with the RepoV plasmid alone or in combination with the PhiC31 (φC31) mRNA (1µgr plasmid: 3µgr mRNA). Flow cytometry analysis of nuEYFP fluorescence in polyclonal iPSC populations obtained by electroporation of RepoV alone or with the PhiC31 mRNA and after for 7 days of G418selection. Fluorescence distributions are shown for the total cell (All cells) and nuEYFP-positive populations (nuEYFP+ cells). Histograms display modal-normalized fluorescence intensity (Log_10_), The red shaded area, delimit the range of nuEYFP negative cells.

### RepoV transfection and integration into human iPS cells

Recombination-mediated enhancer transgenesis in human iPSCs requires co-delivery of the RepoV plasmid and PhiC31 integrase. We first optimized RepoV delivery by electroporation with the Neon Nxt system. For that, we cloned the constitutive human EF1 promoter into RepoV and electroporated the RepoV(EF1p::nuEYFP)^[NeoR]^ plasmid into three independent human iPSC lines (SCTi003, SCTi004, and AG08C5) across a range of electroporation parameters. Twenty-four hours after transfection, we assessed the percentage of nuEYFP+ cells to identify conditions yielding robust and efficient transfection across all lines. This approach identified a common set of electroporation parameters that yielded an average transfection efficiency of 25% of surviving cells (**Fig. 1C**).

We next used these conditions to optimize RepoV and PhiC31 co-transfection (**Fig. 1d** and **Supplementary Fig. S1**). PhiC31 was delivered as in vitro-transcribed mRNA rather than from an expression plasmid to achieve rapid protein production (Zou et al. 2010) and avoiding integration of the PhiC31 vector itself. We therefore tested a range of RepoV amounts and RepoV:PhiC31mRNA ratios. In all cases, 10^6^ iPSCs were electroporated and subjected to 7 days of G418 selection to retain stable transgenic clones (**Supplementary Fig. S1A**) and transgenesis efficiency was scored by the number of NeoR⁺ and nuEYFP⁺ colonies. RepoV transfected alone served as was used a reference, to compare PhiC31-mediated transgenesis with classical plasmid random integration. RepoV alone transfections yielded relatively few NeoR⁺ colonies, with little variation across plasmid doses (**Fig. 1D** and **Supplementary Fig. S1B**)) and, nearly 80% of these colonies lacked detectable nuEYFP expression (NeoR⁺/nuEYFP⁻). We reasoned that these clones reflect random integration events that disrupted the EF1::nuEYFP expression cassette rather than integration into repressive chromatin, since the latter should also silence NeoR. In contrast, co-transfection with PhiC31 mRNA consistently increased the number of stable transgenic colonies over the corresponding RepoV-only conditions, particularly at low plasmid doses, both in absolute NeoR+ colony number and in NeoR+ nuEYFP+ proportions (**Fig. 1D** and **Supplementary Fig. S1B**). This is consistent with PhiC31-mediated recombination integrating the vector without disrupting the reporter cassette. Using the optimal RepoV dosage (1µgr), we then refined the RepoV: PhiC31 mRNA ratio (**Supplementary Figure S1C)** identifying 1:3 as the most efficient condition. Either high RepoV or PhiC31 amounts reduced the number of NeoR⁺ and the proportion of NeoR⁺/nuEYFP⁺ colonies. Besides, NeoR+/nuEYFP− colonies were still detected at every RepoV and PhiC31 dose tested, suggesting that random integration still occurs in the presence of the recombinase.

Variability in transgene activity levels arising from differences in copy number and genomic position effects can strongly hamper the interpretation of candidate sequences in enhancer transgenesis assays. To test whether PhiC31-mediated integration of RepoV reduces this variability, we transfected human iPSCs with RepoV(EF1p::nuEYFP) alone or together with PhiC31 mRNA. After G418 selection of stable transgenic clones, we assessed nuEYFP expression across the polyclonal transgenic population by flow cytometry (**Supplementary Fig. S1A**). As expected, transgenesis with the RepoV vector alone yielded a relatively low proportion of nuEYFP positive cells, amongst which populations expressing high, mid or low nuEYFP levels could be identified **(Fig. 1E)**. Instead, PhiC31-electroporation yielded a considerably higher proportion of nuEYFP+ cells, paralleling the colony assays. Besides, the majority of these cells displayed high fluorescence levels with only minoritarian proportions of low and mid nuEYFP expressing cells. Similar results, were obtained by repeating these experiments using A RepoV plasmid carrying the constitutive CMV promoter (**Supplementary Fig. S1D**). In this case, the proportion of nuEYFP-positive cells was also higher following RepoV/PhiC31 mRNA co-transfection than with RepoV alone. However, two distinct populations of nuEYFP-expressing cells were observed, rather than the single predominant population obtained with the EF1 promoter, suggesting that different CREs may exhibit different degrees of variability in their activity. Nevertheless, the proportion of cells belonging to the low nuEYFP-expression population was again reduced upon PhiC31 co-transfection compared with RepoV alone. Together, these results indicate that PhiC31-mediated integration of the RepoV reporter vector increases transgenesis efficiency while reducing variability in transgene expression. Finally, we evaluated RepoV transgenesis efficiency over candidate sequence ranging from less than 1kb up to 4 kb, observing no correlation among these parameters (**Supplementary Fig S2E**), and indicating that PhiC31 activity is not affected by vector size across the analyzed range.

### Building a versatile RepoV toolkit for enhancer engineering

Beyond enhancer characterization, enhancer transgenesis provides a valuable tool for generating reporter cell and animal lines for lineage tracing, cell morphology /behavior analysis, and population-specific transcriptional and epigenetic profiling (Biase et al. 2018; Espinosa-Medina et al. 2023). To extend the applications of RepoV in organoid cultures, we generated variants carrying different fluorescent reporters and selection cassettes (**Supplementary Fig. S2A-C**).

The original nuEYFP reporter was replaced with EGFP or destabilized TurboGFP (dtTurboGFP), both of which predominantly localize to the cytoplasm and are therefore suitable for monitoring cell morphology and behavior. The shorter half-life of dtTurboGFP enables faster reporter turnover, facilitating the detection of transient enhancer activity. To compare reporter performance, we generated CMV-driven constructs and transiently transfected human iPSCs. Flow cytometry revealed the highest fluorescence levels for RepoV(CMV::nuEYFP) and RepoV(CMV::EGFP), whereas dtTurboGFP showed lower fluorescence intensity, consistent with its shorter half-life (**Supplementary Fig. S2D**). Thus, reporter selection involves a trade-off between signal intensity and temporal resolution, as the lower fluorescence of dtTurboGFP may limit detection of weak enhancers. Additionally, we generated carrying a nuclear Tomato reporter and each reporter configuration was combined with either blasticidin-or puromycin-resistance cassettes to allow generating transgenic lines for multiple reporter cassettes to label distinct cell populations within the same organoid. Together, these modifications expand the versatility of the RepoV platform, allowing reporter configurations to be tailored to different experimental applications.

### Brain organoid cultures

Neurodevelopmental genes are frequently expressed across multiple embryonic brain regions (e.g. (Walther and Gruss 1991; Hatini et al. 1994; Kirsch and Chechik 2016)). Defining the function of their CREs and interpreting disease-associated variants within them, may therefore require assessing their activity across different brain domains. We thus generated cortical (COs), ventral forebrain (VFOs), and retinal organoids (ROs) from iPSCs by adapting published protocols (Bagley et al. 2017; Cowan et al. 2020; Krefft et al. 2018; Lancaster et al. 2017) (**Fig. 2A,B**). We characterized them by histology, immunofluorescence, RT-qPCR, and scRNAseq **(Fig. 2B-E** and **Supplementary Fig. 3,4**).

**Figure 2.**
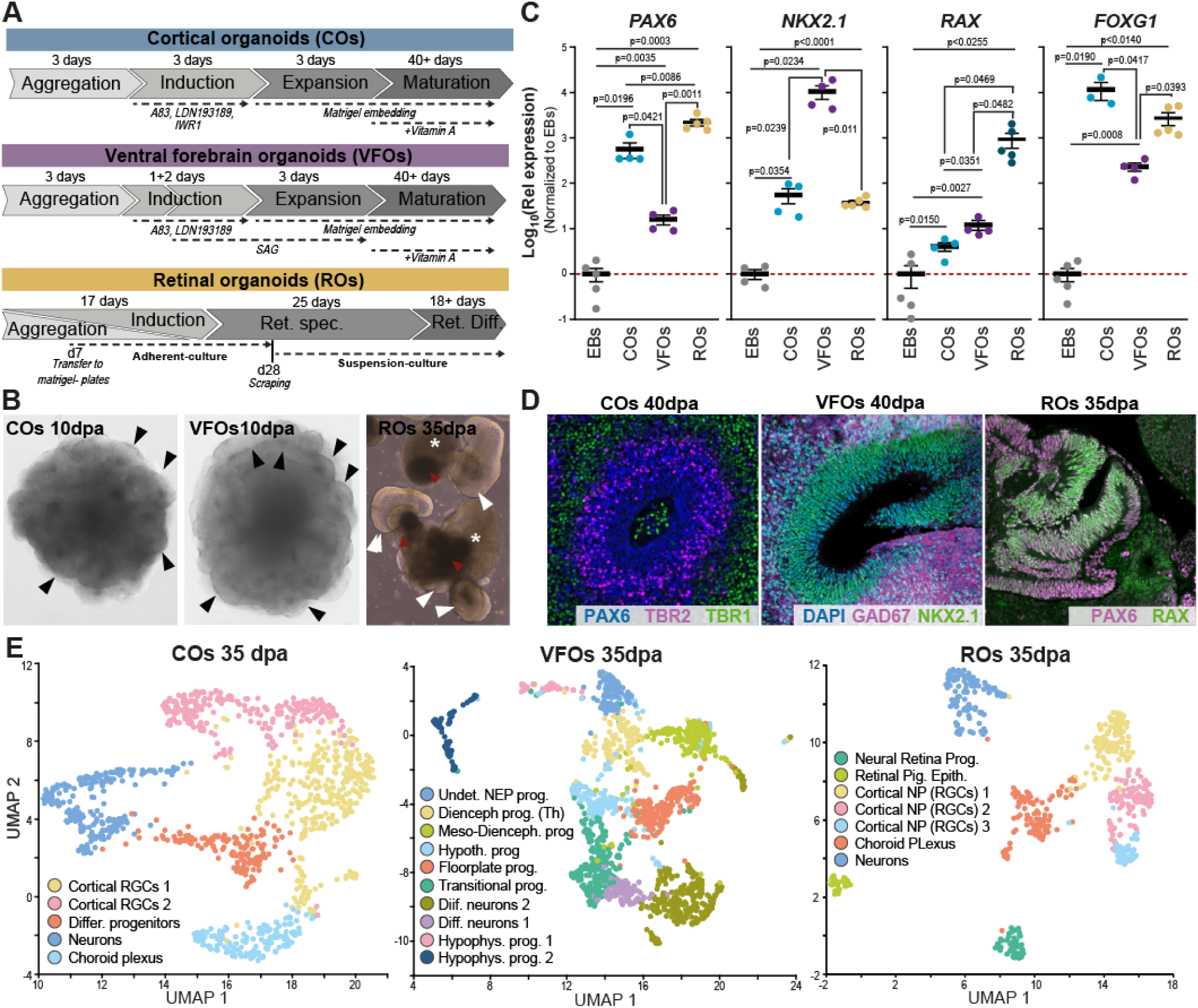
Neural organoid models used in this study. **A.** Scheme depicting the experimental procedures for CO, VFO or RO production. **B.** Bright field microscopy images showing the formation of neurogenic vesicle - like structures in CO-and VFO (black arrowheads) as well as evaginated neuroretina structures (white arrowheads) neighbored by RPE patches (red arrowhead) and abutting from optic dense aggregates (white asterisks). **C.** RT-qPCR experiments showing the enrichment of regional identity markers in mature (40 days) CO, VFOs and ROs compared to undifferentiated EBs. Data are presented as mean ± SEM, with individual data points representing independent samples. Statistical significance is indicated by p-values. **D.** Immunofluorescence images of cryostat sections of CO, VFO and ROs showing the expression of regional identity and cell type-specific markers expected for each organoid, as indicated in the panels. **E.** UMAP plots showing the single-cell transcriptomic profiles of COs, VFOs, and ROs at 35 dpa. Distinct cell populations are identified based on their transcriptional profiles and color-coded according to their annotated identities

A detailed description of organoid formation and characterization is provided in the **Supplementary Information File.** As expected, COs displayed high *PAX6* and *FOXG1* levels compared to undifferentiated embryoid bodies (EBs) while showing only modest *NKX2-1* or *RAX* expression (**Fig. 2C**) and formed organized neurogenic structures containing PAX6+ radial glia, TBR2+ intermediate progenitors and TBR1+/CTIP2+/SATB2+ neurons (**Fig. 2D**). At 35 days post aggregation (dpa), scRNAseq identified cortical radial glia and mainly deep-layer glutamatergic neurons, consistent with an early corticogenic stage (roughly E12.5 in mouse). Instead, ventral forebrain and mesencephalic markers were largely absent in agreement with the qPCR results (**Fig. 2C,E**). Contrary to COs, VFOs contained FOXG1+/NKX2.1+ domains, paralleling the high levels of these markers observed by qPCR, as well as GAD67+ neurons, supporting the presence of ventral forebrain domain and their neuronal progeny (**Fig. 2C,D**). Besides, scRNAseq, analysis revealed progenitors of, hypothalamic, floor plate and diencephalic progenitors, with near absence of dorsal telencephalic markers (**Fig. 2E**). ROs developed neural retina-like structures (PAX6+, RAX+, VSX2+) with adjacent RPE (OTX2+, MITF+) and optic stalk (PAX2+) domains (**Fig. 2B-E**) although, retinal neurons started to be detected only at later stage RO culture stages (56dpa))(**Supplementary Figs. S3C**, **S4F,G**). Interestingly, scRNAseq analysis confirmed these domains but also revealed populations of cortical progenitors and neurons, which, based on immunofluorescence analysis localized mostly in the organoid core, as reported for similar protocols (Gabriel et al. 2021; Liu et al. 2023).

Together, these approaches generated and validate the regional identities of the three organoid models used in the study and support their use for analyzing CREs associated with neurodevelopmental genes in the corresponding embryonic structures.

### PhiC31 transgenesis and organoid cultures for studying human neurodevelopmental enhancers

As a proof of principle that RepoV transgenesis could be applied to evaluate the activity of candidate enhancers we selected evolutionary conserved non-coding sequences previously characterized as active either in the embryonic cortex or in ocular tissues (**Fig. 3**). We first used a CRE located 18.6 kb upstream of the human *EMX1* gene, catalogued in the Vista Enhancer Browser database (Visel et al. 2007) as active in the cortex of E11.5 mouse transgenic embryos (Vista element hs1205, **Fig. 3A,B**) and evolutionary conserved across amniotes. This element inserted into our RepoV(dtTurboGFP) plasmid was used to establish COs from a polyclonal transgenic population. The transgene showed strong activity in these organoids at 20dpa **(Fig. 3C)**, a stage approximately corresponding to the E11.5 mouse embryonic cortex, thus paralleling its reported activity in mice.

**Figure 3.**
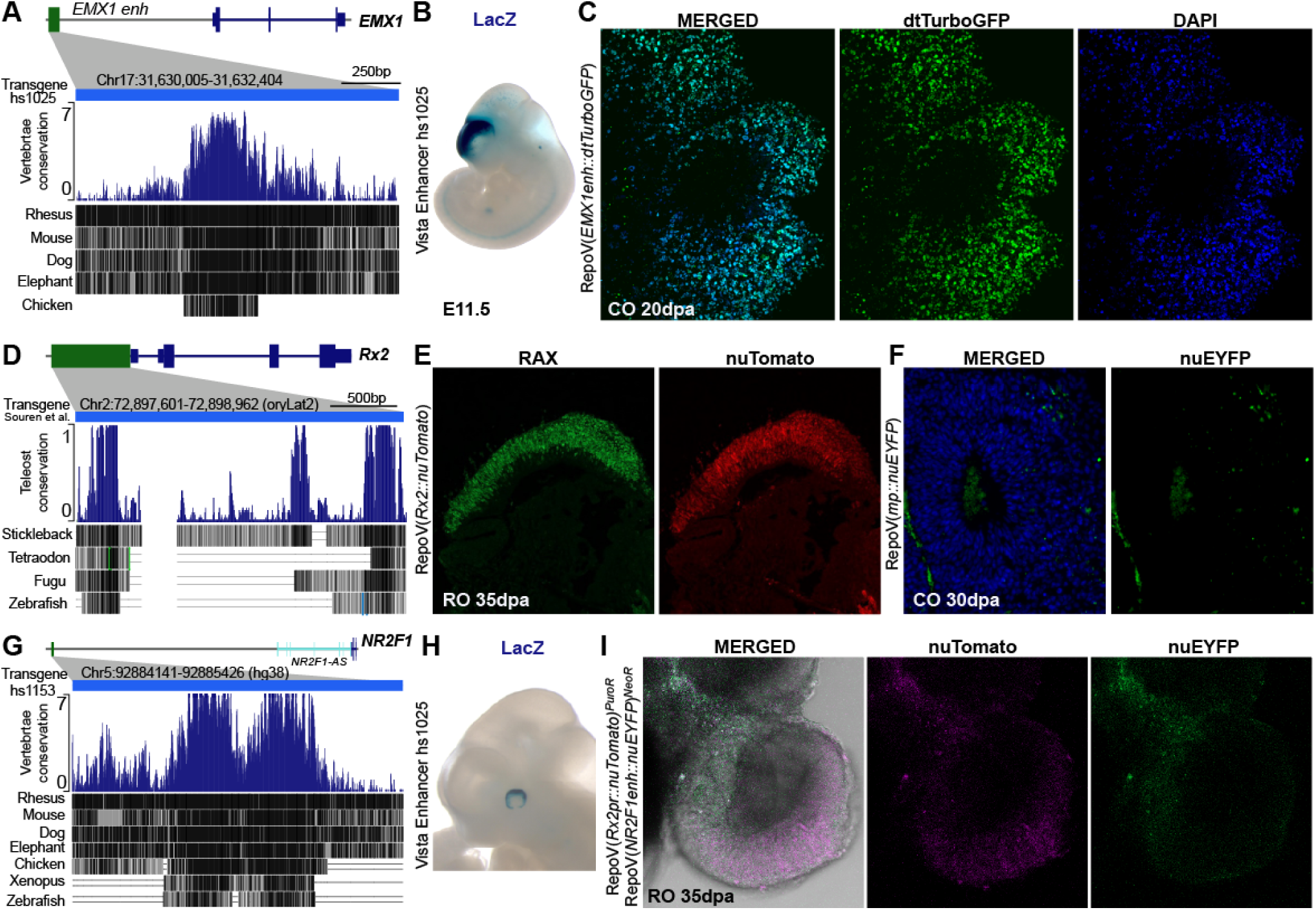
Region-specific human organoids enable the assessment of enhancer activity through RepoV transgenesis. **A, D, G.** Schematic representation of the candidate enhancers associated with *EMX1*, *Rx2*, and *NR2F1*, respectively, showing their genomic location, evolutionary conservation, and previously reported regulatory activity in the developing telencephalon or eye. **B, H.** VISTA enhancer browser images showing LacZ activity driven by the *EMX1* and *NR2F1* CREs in transgenic mouse embryos. **C.** Confocal image of a cortical organoid (CO) derived from a RepoV-transgenic isogenic line, showing activation of the dtTurboGFP reporter. **E.** Confocal image of a retinal organoid (RO) carrying the RepoV(*Rx2p::nuTomato*) construct, showing Tomato expression in the RAX-positive neuroretina. **F.** Representative CO section showing the absence of reporter expression in organoids carrying the empty RepoV(bMP::nuEYFP) construct. **I.** Whole-mount immunofluorescence of an RO derived from a double-transgenic line carrying Repo(*NR2F1*enh::nuEYFP) and Repo(*Rx2p::nuTomato*), showing reporter expression consistent with the respective CRE activity patterns.

A similar approach, but using ROs, was used to functionally evaluate two regulatory elements active in the developing ocular structures (**Fig. 3D-I**). The first was a 2.4 kb upstream promoter region of the medaka *Rx2* gene, which has been extensively validated as an early neuroretinal marker in zebrafish transgenesis and strongly conserved even across distant teleost species (c.a. 350 million years) (**Fig. 3D**, (Souren et al. 2009)). The candidate element was cloned into the RepoV(bMP::nuTomato)^[PuroR]^ plasmid to establish an isogenic iPSC lines. ROs derived from this line showed robust transgene activity in the neuroretinal vesicles but no reporter expression was detected in the CO-like core portion (**Fig. 3E**). The second element was a human non-coding sequence localized 695 kb upstream of the *NR2F1* gene associated to the *NR2F1-AS1* locus (**Fig. 3G**). This element (catalogued as Vista enhancer hs1153) displays robust activity in the dorsal outer portion of the E11 mouse optic cup, possibly corresponding to the presumptive RPE (**Fig. 3 G,H**). This element was cloned into the RepoV(bMP::nuEYFP)^[NeoR]^ vector and then electroporated into the RepoV(Rx2p::Tomato)^[PuroR]^ line. Organoids derived from the resulting polyclonal population showed nuEYFP activity in domains adjacent to the Tomato+ neuroretina vesicle, consistent with the activity reported in the Vista Enhancer browser database.

These experiments demonstrate that RepoV transgenesis can generate combinatorial reporter iPSC lines that enable the simultaneous visualization of distinct cellular populations.

### Applying RepoV transgenesis to characterize the activity of neurodevelopmental enhancers

The vast majority of non-coding sequences identified as potential CREs in the human genome remain functionally uncharacterized. This is particularly challenging for CREs located within exons, as genome editing of their endogenous sequence may affect the gene protein-coding sequence, mRNA stability, or translation. We applied our workflow to analyze the activity of four candidate enhancers identified by epigenetic profiling and associated with the neurodevelopmental genes *POU3F2* and *FOXG1*. The selected candidate sequences displayed chromatin epigenetic landmarks of active CREs and overlap single nucleotide or copy number variants (SNVs/CNVs) associated with cognitive traits or neuropathological conditions (**Fig. 4** and **Fig. 5**).

**Figure 4.**
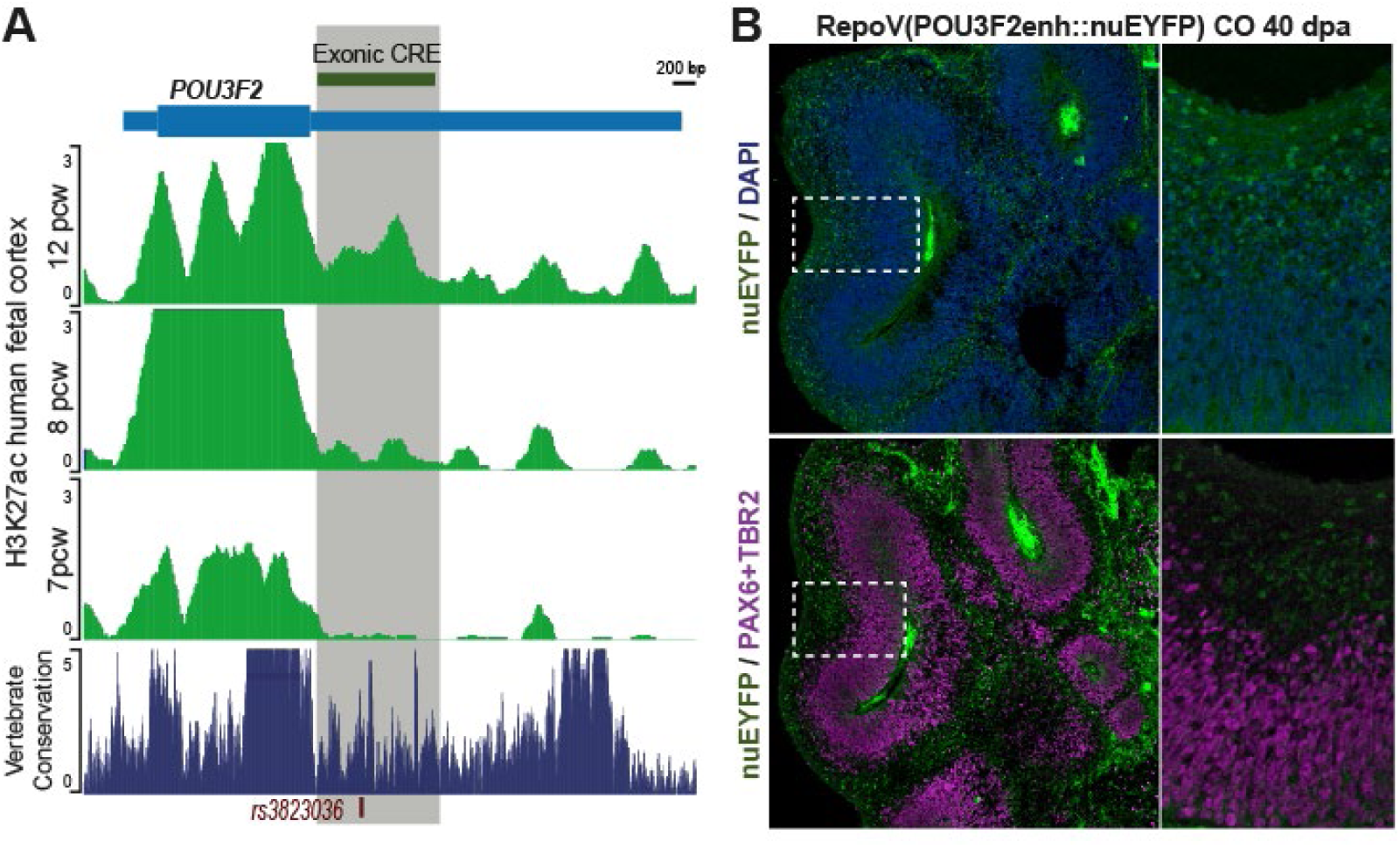
A 3’UTR exonic sequence of the human *PO3UF2* gene displays enhancer activity in human COs. **A.** Schematic representation of the *POU3F2* locus and the candidate CRE identified based on H3K27ac enrichment data from (Reilly et al. 2015)The candidate region overlaps highly conserved vertebrate sequences and shows increased H3K27ac enrichment in late-stage human fetal cortex compared with earlier developmental stages. Besides, a SNV within a conserved region of this element has been associated with human cognitive and behavioral traits. **B, C.** Confocal images of cryostat sections from 40 dpa COs carrying the corresponding RepoV construct, showing nuEYFP reporter expression in cortical plate-like structures. The absence of overlap between nuEYFP-positive nuclei and PAX6-positive radial glial or TBR2+ intermediate progenitors (C) suggests enhancer activity in postmitotic neural populations.

**Figure 5.**
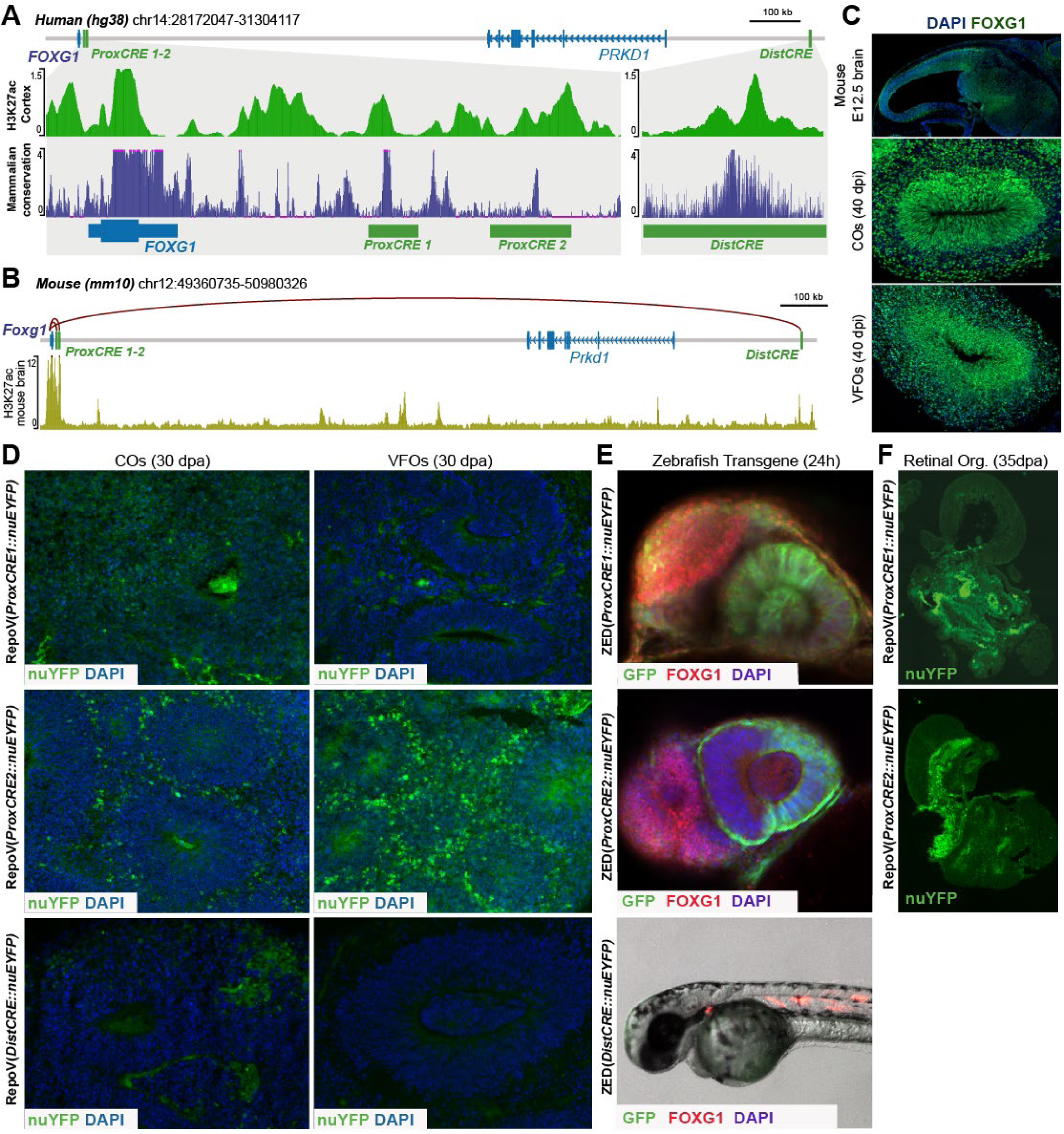
Characterization of CREs associated with the human *FOXG1* gene. **A,B.** Schematic representation of the human and mouse *FOXG1* loci, showing the location and evolutionary conservation of candidate CREs, H3K27ac enrichment in human fetal cortex (Reilly et al. 2015) or mouse brain (ENCODE4 dataset) as well as the CHIPAT contacts among the mouse ortholog elements and the *Foxg1* promoter (Bertolini et al. 2019). **C.** Confocal images of E12.5 mouse forebrain and 40 dpa cortical (COs) and ventral forebrain organoids (VFOs), showing FOXG1 expression in dorsal and ventral pallial regions and their organoid counterparts. **D.** Microscopy images of cryostat sections from COs and VFOs transgenic for each candidate CRE reporter constructs. **E.** icroscopy and stereomicroscopy images of F1 zebrafish embryos transgenic for the proximal FOXG1 CRE1 and CRE2 constructs, showing EGFP reporter expression and FOXG1 immunostaining. No EGFP expression was detected in zebrafish F0 or stable lines carrying the distal FOXG1 CRE, despite activation of the control Tomato reporter in larval musculature. **F.** Microscopy images from cryostat sections of COs and VFOs transgenic for the candidate CRE’s constructs.

The putative *POU3F2* regulatory sequence overlaps the 3’ UTR portion of the locus, is evolutionary conserved across mammals and displays stage specific enrichment of the H3K27ac enhancer epigenetic mark in human fetal cortex (**Fig. 4A**). A SNV in this element has been associated with increased mathematical capabilities, and attention deficit disorder in GWAS association studies (**Fig. 4A** rs3823036; (Lee et al. 2018; Rao et al. 2022)). COs generated from the corresponding polyclonal RepoV(POU3F2enh::nuEYFP) transgenic iPSC population displayed nuEYFP expression into their cortical plate-like structures **(Fig. 4B**), but not in their ventricular/ subvetricular regions (immunostained for PAX6/TBR2) suggesting that the transgene is activate in post-mitotic neural precursors and/or differentiating neurons. This strongly supports the enhancer function of this exonic 3’UTR sequence, and suggests that alteration of its enhancer activity by the rs3823036 SNV could contribute to the neurobehavioral-traits-described in patients.

The three additional candidate CREs were associated with the locus of the human *FOXG1* (**Fig. 5A**), a gene expressed in dorsal and ventral telencephalic domains and in the developing eye (Hatini et al. 1994). Two proximal elements lie 11.3 and 15 kb downstream of the *FOXG1* TSS and were designated ProxCRE1 and ProxCRE2, respectively, while the third is located approximately 1.5 Mb downstream of the locus (DistCRE). All three elements display strong evolutionary conservation across mammals and H3K27ac enrichment in the human embryonic cortex (**Fig. 5A**). Analysis of ChIA-PET data (Bertolini et al. 2019; D’Aurizio et al. 2022) revealed that their mouse orthologues contact the *Foxg1* promoter and are bound by SOX2, a key neurodevelopmental transcription factor involved in telencephalic specification and cortical neurogenesis (**Fig. 5B**). Importantly, CNVs encompassing these elements are associated with neurodevelopmental delay and Rett-like syndrome patients (D’Aurizio et al. 2022) as well as with Autism Spectrum Disorder (unpublished data).

Each human sequence was cloned into the RepoV(bMP::nuEYFP) vector to generate polyclonal transgenic iPSC lines that were used to generated both COs and VFOs, to cover both *FOXG1* telencephalic expression domains (**Fig. 5C** and **Supplementary Figs. S3**). RepoV(ProxCRE1::nuEYFP) displayed modest but reproducible activity in COs, but not VFOs, whereas RepoV(ProxCRE2::nuEYFP) was active in both. **(Fig. 5D**). Besides the localization of the nuEYFP+ cells suggest that both ProxCRE1 and ProxCRE2 elements are active in potst mitotic neural precursors / neurons. The DistCRE cassette instead failed to drive reporter expression in either organoid models (**Fig. 5D**). To further assess their regulatory potential, the three elements were cloned into the ZED vector and analyzed in F0 and F1 transgenic zebrafish embryos (**Fig. 5E**). Consistent with the organoid results, ZED (DistCRE::EGFP) embryos showed no EGFP expression despite effective transgenesis (positive tdTomato expression in muscle), whereas ProxCRE1 and ProxCRE2 drove strong EGFP expression in the optic cup of 24-hpf embryos, but little or no expression in the telencephalon, apart from a small group of EGFP+ cells in ProxCRE1 transgenics. Notably, ProxCRE1 and ProxCRE2 did not drive nuEYFP expression in VSX2+ retinal structures present in human ROs (**Fig. 5F**), although ProxCRE1 produced patches of nuEYFP+ nuclei in the inner organoid portion, within the VSX2 negative domain, and consistent with its activity in COs.

Collectively, these results indicate that ProxCRE1 and ProxCRE2 contain regulatory information that differs between zebrafish and human organoid models. Such discrepancies may reflect evolutionary changes in the trans-regulatory networks controlling embryonic structure development as postulated in other studies (Ariza-Cosano et al. 2012). The lack of reporter expression associated with DistCRE instead suggests that its functions may depend on cooperation with other CREs or additional chromatin elements, or involve non-canonical enhancer activities.

Overall, these findings demonstrate that combining PhiC31-mediated transgenesis in human iPSCs with organoid cultures provides a useful approach for evaluating candidate regulatory sequences in relevant human neurodevelopmental contexts.

## DISCUSSION

Here we describe RepoV, a PhiC31 integrase-based vector that drives targeted, predominantly single-copy integration of insulated enhancer-reporter cassettes into endogenous pseudo-attP sites of the human genome. Combined with region-specific neural organoids, RepoV trangenesis allowed us to recapitulate the reported activity of three previously characterized regulatory elements, and to demonstrate the enhancer functions of an exonic CRE of *POU3F2*, of two *FOXG1* proximal CREs. Differences in the reporter activity of FOXG1 enhancers between human organoids and zebrafish embryos suggest that some CREs may exhibit species-dependent activity, as previously reported (Ariza-Cosano et al. 2012a). Besides, the organoid results are more consistent with FOXG1 expression and the pathological phenotypes associated with disease-related regulatory variants identified in patients, highlighting the importance of human-based developmental models for evaluating human CREs. Together, these findings establish a workflow for assessing candidate human regulatory sequences at cellular resolution and within a human developmental context, advancing current approaches to functional CRE characterization.

### Enhancer transgenesis in organoid models complements existing approaches

Functional characterization of the human regulatory genome remains a major obstacle to understanding coordinated gene expression and how non-coding variants contribute to disease. This task is compounded by the combinatorial nature of gene regulation, in which multiple elements can act additively, redundantly, or synergistically. Computational approaches integrating bulk and single-cell epigenomic and transcriptomic data can prioritize candidate CRE and disease-associated variants (Sánchez-Gaya and Rada-Iglesias 2023; Schubach et al. 2024), but functional assignment requires complementary experimental strategies. Reporter assays test whether individual CREs or combinations thereof are sufficient to activate or repress transcription, whereas genome editing tests whether they are necessary in their native context.

Human brain organoids provide a relevant model for investigating regulatory mechanisms during neurodevelopment, allowing to exploit human iPSC amenability for transgenesis and genome editing manipulations. In fact, lentivirus-based massively parallel reporter assays (MPRAs) have been used to assess the activity of numerous candidate enhancers and the effects of putative regulatory variants in these models (Capauto et al. 2024; Deng et al. 2024; Noack et al. 2023). STARRseq is a powerful high throughput assay for large scale evaluation of enhancer activity in cell lines or primary cultures, including human neural progenitors that has to provided key insights on the regulatory logic of transcription factor binding sites and enhancer promoter compatibility (Bergman et al. 2022; Gaynor-Gillett et al. 2025; Zabidi et al. 2015). However, these assays are hampered by limitations in the size of the candidate elements analysed and/ or the possibility of evaluating combination of them. Besides, their sensitivity to analyse activity of weak enhancers or elements active in rare cell population is subject to limitations of bulk or single cell sequencing approaches (Preissl et al. 2023).

The PhiC31-mediated enhancer transgenesis method described here provide a simple and efficient approach to evaluate the activity of candidate regulatory sequence in human organoid cultures, overcoming limitations of animals models in terms of evolutionary divergence of the embryonic structure or organ being analysed while fulfilling 3R principal of animal research (Hubrecht and Carter 2019). Our results indicate that RepoV transgenesis efficiency is not constrained by the size of the regulatory element cloned and therefore allows testing relatively large candidate sequences (in this study we evaluated elements of up to 3.5kb) or combinations of elements. Thus, our method is particularly suited for the in-depth characterization of defined sets of human candidate regulatory sequences (or combinations thereof), enabling the assessment of their activity with cellular resolution, offering an alternative approach to the mentioned high throughput strategies and complementing animal-based enhancer transgenesis. Finally, this method is well suited for establishing enhancer-reporter cell lines that enable the selective labeling of specific cell populations within organoid cultures, facilitating their subsequent transcriptional, epigenetic or cell behavior characterization. In fact, the RepoV toolkit developed in this study supports the integration of multiple reporter cassettes carrying distinct fluorescent markers, with either nuclear or cytoplasmic localization, thereby enabling the simultaneous labeling of different cell populations and accommodating a broad range of experimental needs.

### Limitations and further development of the RepoV transgenesis

Our PhiC31 mediated transgenesis approach is designed to direct RepoV plasmid integration to endogenous pseudo attP sites of the human genome which had been previously identified by (Chalberg et al. 2006b). This work demonstrated that integration events predominantly occur in e restricted subset of these site, which likely contribute to yield more robust and consistent expression levels compared to random plasmid integration. This strategy is meant to simplify application of this transgenesis approach to virtually any human iPSC or ESC line without the requirement of engineer attP landing sites at specific locations. However, this variable landing site integration partially hampers direct comparison of allelic versions of a candidate regulatory sequence, for example for the assessment of how SNVs or small CNVs impact enhancer activity. Engineering of attP landing sites for the PhiC31 integrase or other serine recombinases at characterized safe harbor loci of the human genome would overcome this limitation, yet at the expense of limiting transgenesis assays to previously edited iPSC/ ESC lines. Thus, balancing assay versatility and specificity depends on considerations related to the biological question being addressed.

Another aspect requiring consideration, is that while organoid cultures provide an interesting platform for the evaluation of human CRE activity, the lack of anatomically defined developing structure impose constrains for the identification of the specific cell types displaying reporter expression, imposing the need of specific markers for the populations of interest. This is particularly true for organoids recapitulating the development of different brain domains, as exemplified by our VFOs, which include different ventral telencephalic, hypothalamic and diencephalic neural progenitors and neurons. However, the improvements and progressive cost reduction of scRNAseq approaches allow to envisage the application of this type of analysis for the characterization of enhancer activity, as already performed in some MPRAs.

Finally, while enhancers constitute the main functional category of CREs, these include other types of elements such as silencers, insulators and tethering elements. While comparatively understudied compared to enhancers, silencers have been shown to importantly contribute to gene regulation and that mutations in these elements also relate to pathological conditions (Segert et al. 2021; Doni Jayavelu et al. 2020). The RepoV constructs generated for this study allow characterizing enhancer activity of candidate CREs while minimizing confounding genome position effects. However, the architecture of the reporter cassette, paralleling that of other classical transgenesis vectors strongly biases its applicability towards the analysis of enhancers rather than other type of CREs. However, with modifications to the RepoV vector inspired by works characterizing the activity of silencers (e.g.(Pang and Snyder 2020)), the same methodological workflow could be applied to the analysis of these CREs, thus further expanding the potential of recombinase-mediated transgenesis and organoid cultures for the study of the regulatory genome.

## MATHERIAL AND METHODS

### iPSC cell and organoid cultures

The established iPSC lines AG08C5 (Badja et al. 2014), SCTi003A and SCTi004A (StemCell Technologies, #200-0511 and #200-0769) were used for this study. All cell lines were verified negative for the most common chromosomal anomalies using the hPSC Genetic Analysis Kit (StemCell Technologies). In all cases, cells were cultured using eTeSR media (Stem Cell Technologies) in cell culture-treated flasks coated with Geltrex (Gibco). Cells were passed at aprox 70% confluence dissociating them to clusters with the Gentle Cell Dissociation Reagent (GCDR; StemCell Technologies).

### Plasmid construction

The RepoV vector was generated by assembling fragments synthetized in vitro or PCR amplified from different plasmids using the NEBuilder HiFi DNA Assembly Cloning Kit (NEB). For the in vitro transcription of the PhiC31 integrase mRNA the coding sequence of the human optimized PhiC31 gene was PCR amplified from the pET-PhiC31o-pA plasmid (Tasic et al. 2011) (a gift from Liqun Luo, Addgene plasmid #52491) and cloned into the pcDNA 3.1 A(-) vector (Thermo Fisher). PhiC31 mRNA was synthetized with HiScribe T7 mRNA Kit with CleanCap Reagent AG (NEB). Similarly, the *CMV* and *EF1* promoters and the *EMX1*, *Rx2*, *NR2F1* and *POU3F2* CREs were PCR amplified and either directly cloned into the RepoV vector by TA cloning or NEB builder HiFi assembly. The *FOXG1* candidate *CREs* were PCR and initially cloned into the PCR 8 GW entry vector (Invitrogen) and then transferred to either the RepoV or ZED vectors either via LR clonase II reaction or by NEB builder assembly

### Electroporation of human iPSCs and transgenic clone selection

The Neon Nxt electroporation system was used for human iPSC electroporation. Briefly, cells were dissociated to single cells using the GCDR, and washed in PBS supplemented with 10µM of the Y27632 ROCK inhibitor (StemCell Technologies). After centrifugation the cell pellet was resuspended in 200-500ul of electroporation R buffer to a final concentration of 10^7^ cells/ml. Either 10 or 100μL of this cell suspension were used for electroporation with the Neon NxT Electroporation System 10μL or 100μL Kit (Invitrogen), according to manufacturer instructions. Optimization of electroporation parameters was performed following the pre-programmed Neon NxT 24-well optimization protocols. Optimal parameters were identified as 2 pulses of 20ms at 1200V input current. After electroporation, cells were plated on Geltrex coated flasks or 6 well plates in 2.5ml of eTeSR supplemented with 10µM of Y27632. After 24h post-electroporation, the media was replaced with fresh eTeSR and allowed to recover for additional 48h-72h with daily media replacement. To select cells carrying stably integrated transgenes, electroporated cells were cultured for 5–7 days in eTeSR supplemented with either 200µg/mL G418 (Merck) or 200 ng/mL puromycin (Merck), depending on the RepoV plasmid used, with daily medium changes. Antibiotic concentrations were set at the minimum doses required to completely eliminate mock-electroporated iPSCs within 5 days. Following selection, surviving cells were allowed to recover for approximately 3–4 days before being used for passaging, cryopreservation, or isogenic clone isolation.

### Flow cytometry analysis and FACS-mediated isolation of iPSC clones

PSCs were dissociated, washed, and resuspended in FAC buffer supplemented with 10 µM Y-27632. Dead cells were excluded by propidium iodide staining, and transgene expression was analyzed using a BD FACSCanto cytometer, with mock-electroporated cells as negative controls. Isogenic clones were obtained by FACS according to (Sarmah et al. 2026) with minor modifications (Supplementary information file and using a FACSAria Fusion (100 µm nozzle, 20 psi, 20–30 µL/min). Cells were sorted into Geltrex-coated 96-well plates containing eTeSR supplemented with CloneR2. After approximately 10 days, clones were dissociated to be expanded and a fraction of the cells were used for genotyping by PCR using RepoV-specific primers. A fraction of each polyclonal population was cryopreserved in Synth-a-Freeze medium.

### Organoid cultures

COs were generated following established protocols (Krefft et al. 2018; Lancaster et al. 2017) with minor modifications (**Fig. 2A**). Briefly, iPSCs were dissociated and seeded (8,000 cells/well) in U-bottom, low-adhesion 96-well plates in eTeSR containing 20 µM Y-27632 to promote EB formation. At 450–550 µm, EBs were transferred to neural induction medium containing LDN193189 (0.2 µM), A83-01 (0.5 µM), and IWR1 (3 µM). After 3 days, organoids were embedded in Geltrex and cultured in expansion medium on an orbital shaker (37°C, 5% CO₂). Three days later, maturation medium was introduced to promote neuronal differentiation. VFOs were generated similarly, except that induction began with 24h ininduction mediu containing LDN193189 and A83-01, followed by the addition of SAG (0.1 µM) to both induction and expansion media.

Retinal organoids were generated according to Cowan et al. (2020). Dissociated iPSCs were seeded in agarose microwell arrays, and aggregation medium was gradually replaced with neural induction medium through daily partial medium changes. At 7 days post-aggregation (dpa), aggregates were transferred to Geltrex-coated 6-well plates. Retinal differentiation was induced at 17 dpa, and at 28– 30 dpa, organoids were transferred to suspension culture on an orbital shaker.

Detailed medium compositions are protocols and media composition are provided in the Supplementary Information file.

### Histological processing and immunofluorescence analysis

Organoids were fixed in 4% PFA overnight at 4°C, cryoprotected in 30% sucrose, embedded in 1:1 gelatin/sucrose, frozen in isopentane (−40°C), and cryosectioned (15 µm). Sections were permeabilized, blocked, and incubated with primary antibodies overnight, followed by secondary antibodies for 2 h. Antigen retrieval was performed when required. Images were acquired using a Leica Stellaris 8 confocal microscope with a 20×/0.75 objective and 2–5 µm optical section intervals.

### RNA extraction and real time PCR

RNA from organoids was extracted using the NucleoSpin RNA Purification Kit (Macherey-Nagel). Reverse transcription and qPCR were performed using the High-Capacity RNA-to-cDNA kit (Applied Biosystems) and the Fast Syber Green Master Mix (Applied Biosystems) respectively.. Gene expression levels were normalized to the housekeeping gene HMBS.

### Zebrafish transgenesis

For each *FOXG1* enhancer candidate, one-cell-stage zebrafish embryos were injected as previously described (Bessa et al. 2009). Enhancer activity was assessed in F0 larvae at 24 and 48 hours post-fertilization (hpf) based on GFP fluorescence. Transgenesis efficiency was evaluated at 48 hpf by detecting the Tomato reporter expression in the axial musculature. The Tomato + larvae were raised to adulthood and crossed with wild-type zebrafish to establish stable transgenic lines for further validation of enhancer activity.

### Single cell RNAseq experiments

For scRNA-seq experiments, organoids were dissociated into single cells following Kanton et al. (2019), with minor modifications. Briefly, organoids were incubated in a papain dissociation solution at 37°C for 45 min with agitation and dissociated by gently pipetting. Dissociation was stopped by adding 10 mL of stopping solution (HBSS supplemented with 1% BSA, 1× Complete Protease Inhibitor Cocktail (Roche), and 10 mg/mL ovomucoid inhibitor (Worthington), followed by centrifugation at 200 × *g* for 5 min. The cell pellet was washed twice with stopping solution and resuspended in the Illumina resuspension buffer. Only samples with viability above 90% were processed further. Cell barcoding and library preparation were performed using the Illumina Single Cell 3′ RNA Prep, T2 by the Experimental Genomics Core Facility of the CBM, using approximately 5,000 cells per sample as starting input. Libraries were sequenced using 100-bp paired-end reads on an MGI DNBSEQ instrument.

### Data analysis

Raw FASTQ files generated from scRNAseq experiments were processed using the PipeSeeker pipeline, with sequencing reads aligned to the human GRCh38/hg38 reference genome. Following processing, the resulting gene expression data were imported into the Trailmaker platform (Parse Biosciences) for downstream analysis. Quality control procedures were performed to assess data quality and identify low-quality cells. The processed data were then used for unsupervised clustering and UMAP representation based on the Seurat package. Cell clusters were manually annotated based on the expression of known cell-type-specific marker genes.

H3K27ac ChIP-seq data for the human fetal cortex from (Reilly et al. 2015) were downloaded from GEO (GSE63648) and converted to GRCh38/hg38 coordinates using the UCSC Genome Browser LiftOver tool. Statistical analyses and qPCR data visualization were performed using GraphPad Prism 9.0. Artificial intelligence (AI) was used solely for English-language editing and text refinement.

### Data, plasmid and protocol availability

The RepoV vectors used in this study are available upon request. Detailed, step-by-step protocol for iPSC cell electroporation and selection are provided in the Supplementary Information File. Single cell RNAseq data are deposited in GEO.

## ACKNOWLEDGMENTS

We thank the Experimental Genomics, Cytometry, and Microscopy Facilities for their support with scRNA-seq library preparation, genetic analyses, flow cytometry, clone isolation, and imaging. We are grateful to Giorgia Pozzolini, Elena Bacchelli, and members of the SNK laboratory for fruitful discussions. This work was supported by the Spanish Ministry of Science and Innovation (MCIN) and the State Research Agency (AEI) through grants CNS2023-144348 (Plan de Consolidación Investigadora), co-funded by the European Union through NextGeneration EU/PRTR, and PID2021-127859NA-I00 (Plan Estatal de Investigación Científica y Técnica y de Innovación), co-funded by the European Regional Development Fund (FEDER; “A way of making Europe”). LB was supported by a Ramón y Cajal fellowship (RYC2020-030257-I), funded by MCIN/AEI. EPS was supported by a predoctoral fellowship (PREP2022-000697) funded by MCIN, AEI, and the European Social Fund Plus (ESF+), associated with project PID2022-16831OB-100. JMG was supported by a predoctoral fellowship (PRE2022-101884) associated with the Severo Ochoa Excellence Accreditation of the Centro de Biología Molecular Severo Ochoa (CBM, CSIC-UAM; CEX2021-001154-S), funded by the Spanish State Research Agency (AEI). SKN and EZ were supported by Fondazione Telethon– Fondazione Cariplo (GJC21176), the European Union (ERANET-NEURON Brain4Sight; NEURON_NDD-255), and the Italian Ministry of University and Research (PRIN-PNRR 2022JKJJ4F). PB, acknowledge support from the EU Brain4Sight ERANET-NEURON project (NEURON_NDD-255) and PCI2022-132938.

## AUTHOR CONTRIBUTIONS

JMG: RepoV construct generation and transgenesis optimization, EMX1 enhancer transgenic generation and organoid analysis, CO culture protocol optimization., DMS: RepoV construct generation and transgenesis optimization, VFO culture protocol implementation and *POU3F2* and *FOXG1* enhancer transgenesis. EPS: *Rx2* and *NR2F1* CRE construct cloning and trangenesis, RO culture implementation. RMF: RepoV transgenesis optimization, organoid culture, and histological and immunofluorescence processing of *FOXG1* CRE transgenic organoids. NT: ZED plasmid cloning and zebrafish transgenesis. LB conceived and designed the transgenesis and organoid culture strategies, coordinated the project, and wrote the manuscript. PB. conceived and designed the zebrafish transgenesis experiments and contributed to manuscript writing and revision. SNK contributed to the design of *FOXG1* enhancer experiments and to manuscript writing and revision.

