## Supplementary FIgures for "ORGANOID-BASED ASSESSMENT OF ENHANCER ACTIVITY VIA RECOMBINATION-MEDIATED TRANGENESIS"

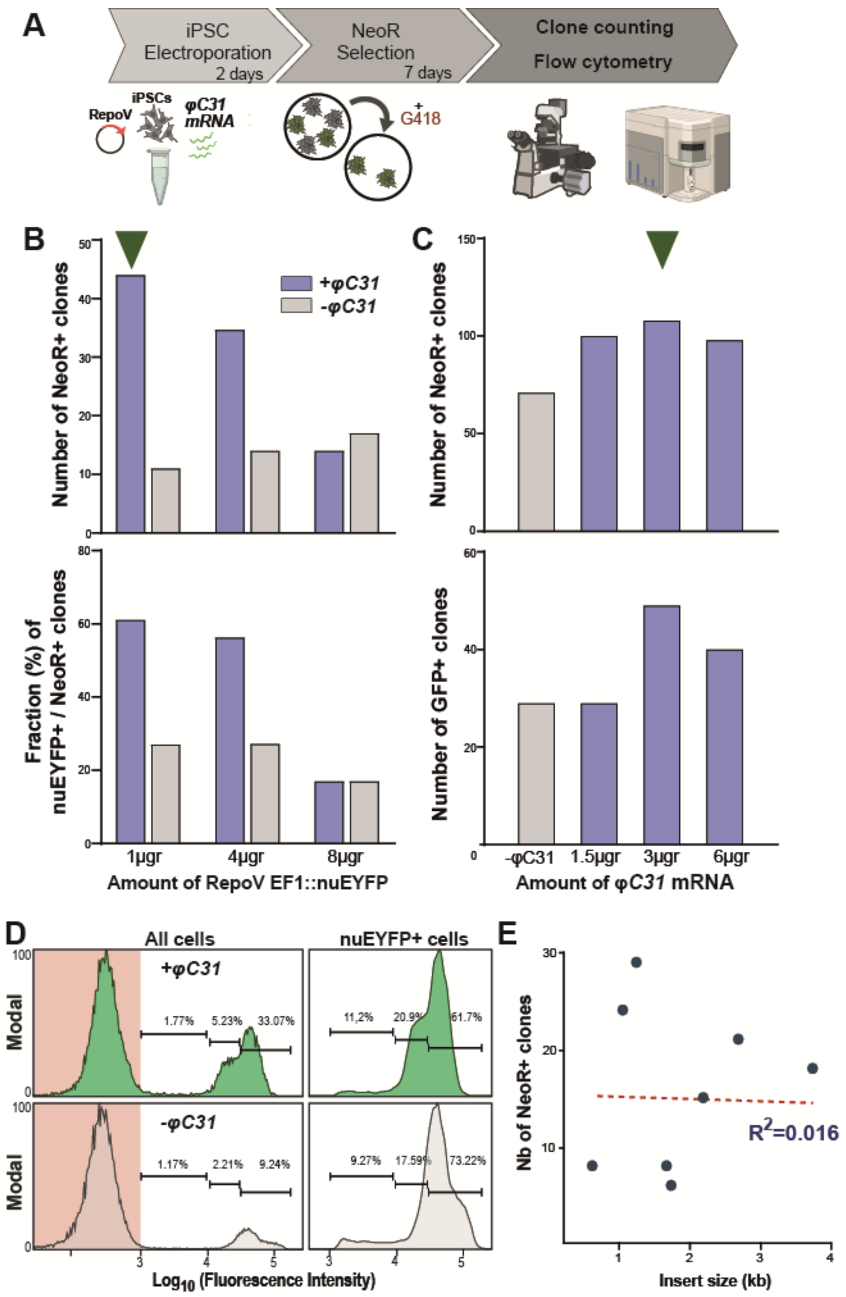


**Supplementary Figure S1. Optimization of RepoV transgenesis. A.** Scheme of the methodological workflow for analyzing transgenesis efficiency upon RepoV electroporation in human iPSC. **B, C.** Bar graphs showing the number of NeoR+ clones and the proportion of the nuEYfP+ ones after G418 selection. We assessed different RepoV amounts (B) and different RepoV: PhiC31 mRNA ratios (C). **D.** Flow cytometry analysis of transgene expression level in a polyclonal iPSC population transgenic for the RepoV(CMV::nuEYFP) reporter construct. **E.** Scatter dot plot representing the transgenesis efficiency (number of NeoR+ clones obtained per electroporation) of RepoV constructs with different insert size and showing the lack of correlation among these variables.

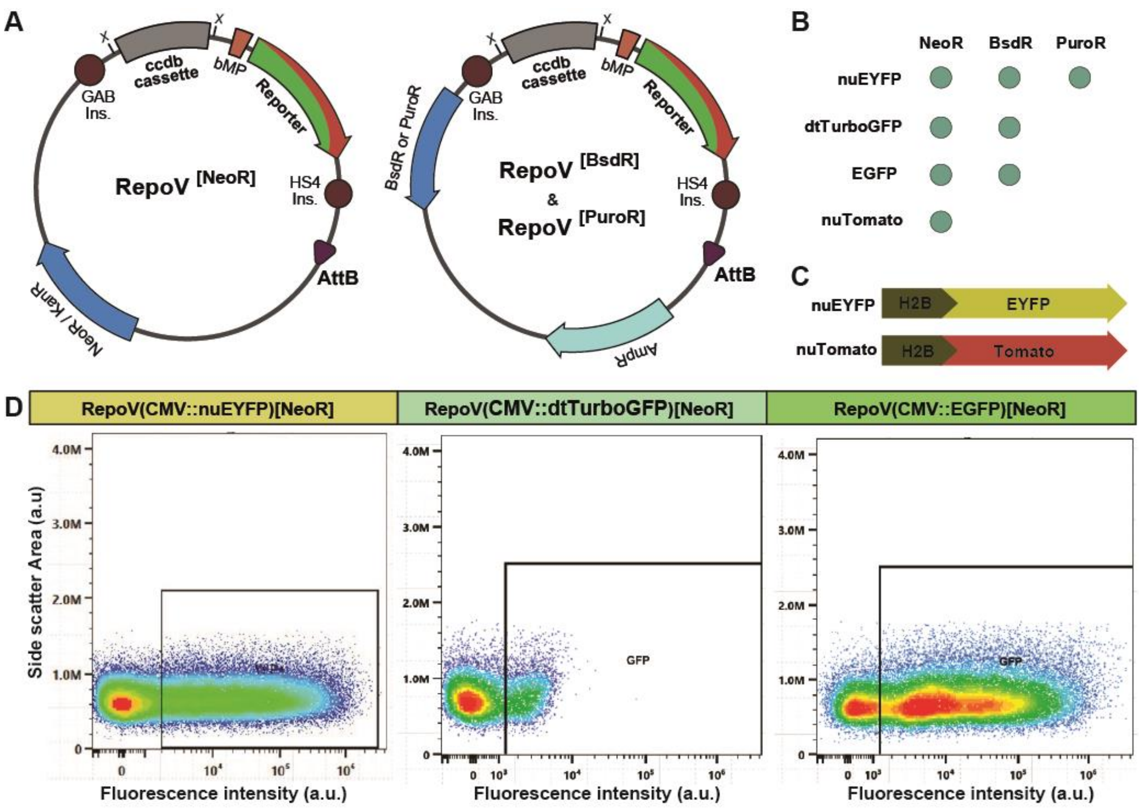


**Supplementary Figure S2. The RepoV transgenesis toolkit. A.** Scheme of the different RepoV backbone vectors generated in this study. **B.** Summary scheme of the different resistance and reporter cassettes used in the RepoV vectors. **C.** The unclear version of the EYFP, cherry and Tomato reporters were generated by fusing their respective coding sequences at the C terminus of the H2B histone coding sequence. **D.** Flow cytometry comparison of the nuEYFP, EGFP and dtTurboGFP expression levels.

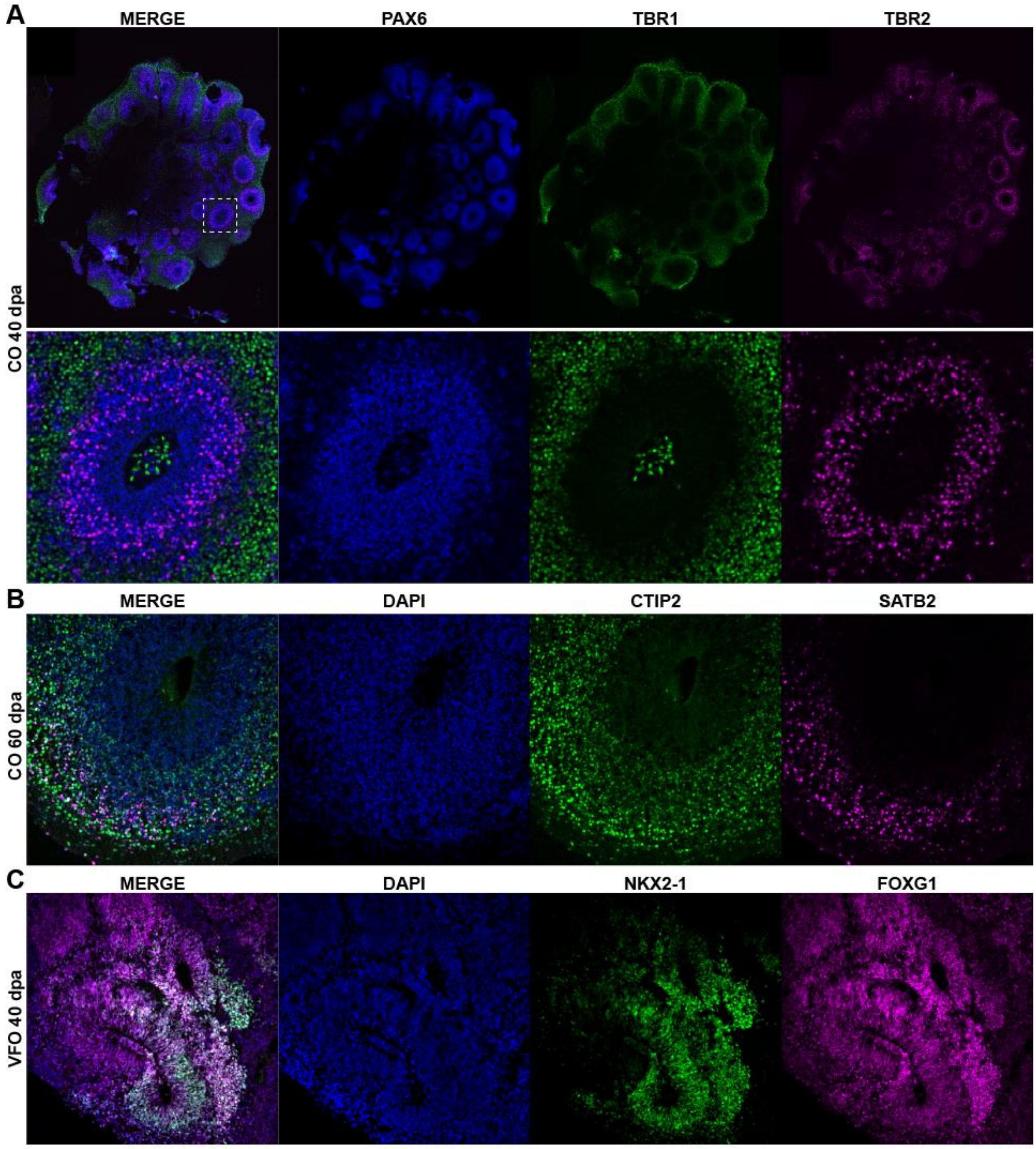


**Supplementary Figure S3. Regional and cellular marker of COs and VFOs. A-D**. Fluorescence microscopy images of cryostat sections from COs (A,B) and VFO (C,D) stained for the indicated markers. The vesicle delineated in a white dashed rectangles the top A panel is shown at higher magnification in the bottom panels

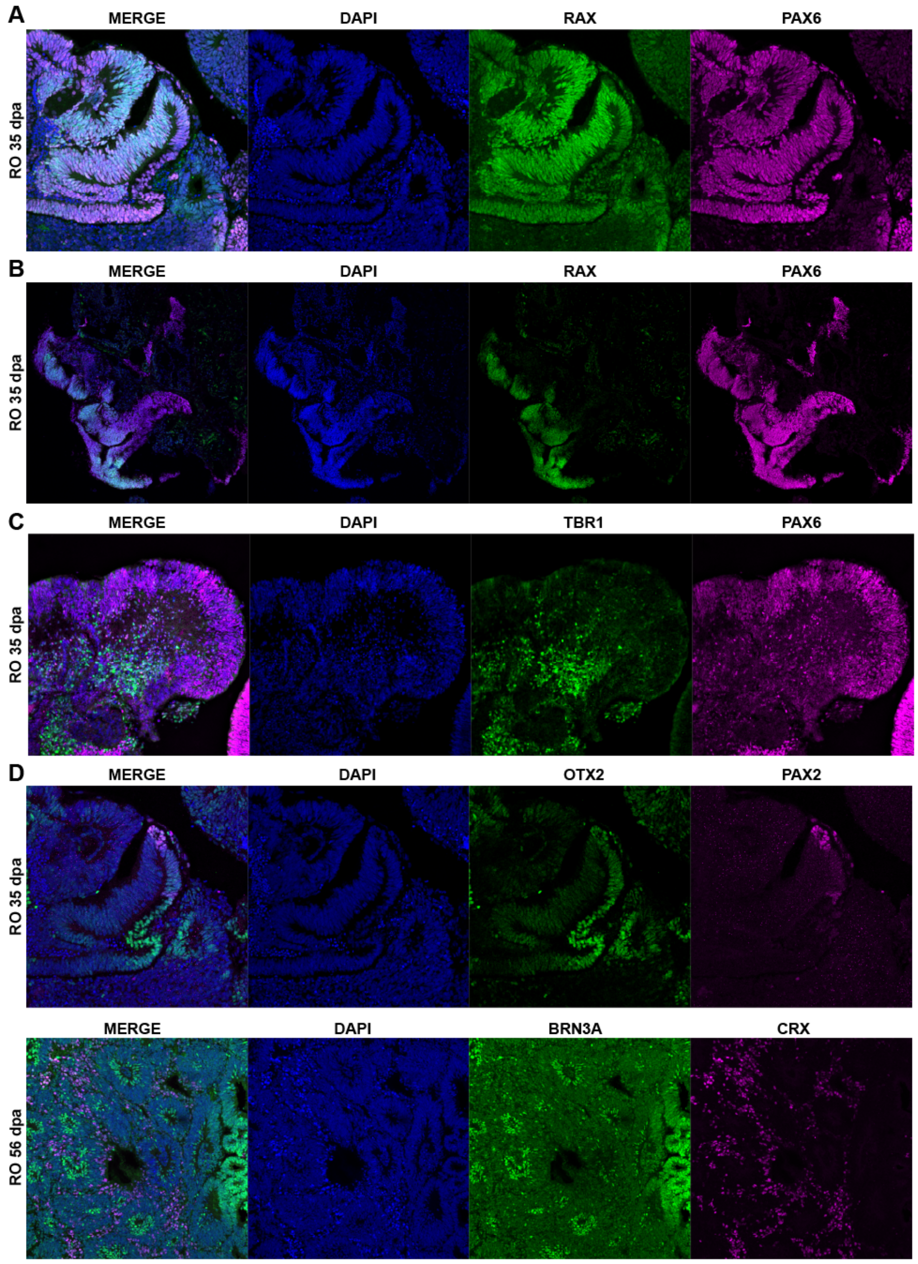


**Supplementary Figure S4. Regional and cellular marker of ROs. A-E**. Fluorescence microscopy images of cryostat sections from ROs stained for the indicated markers.

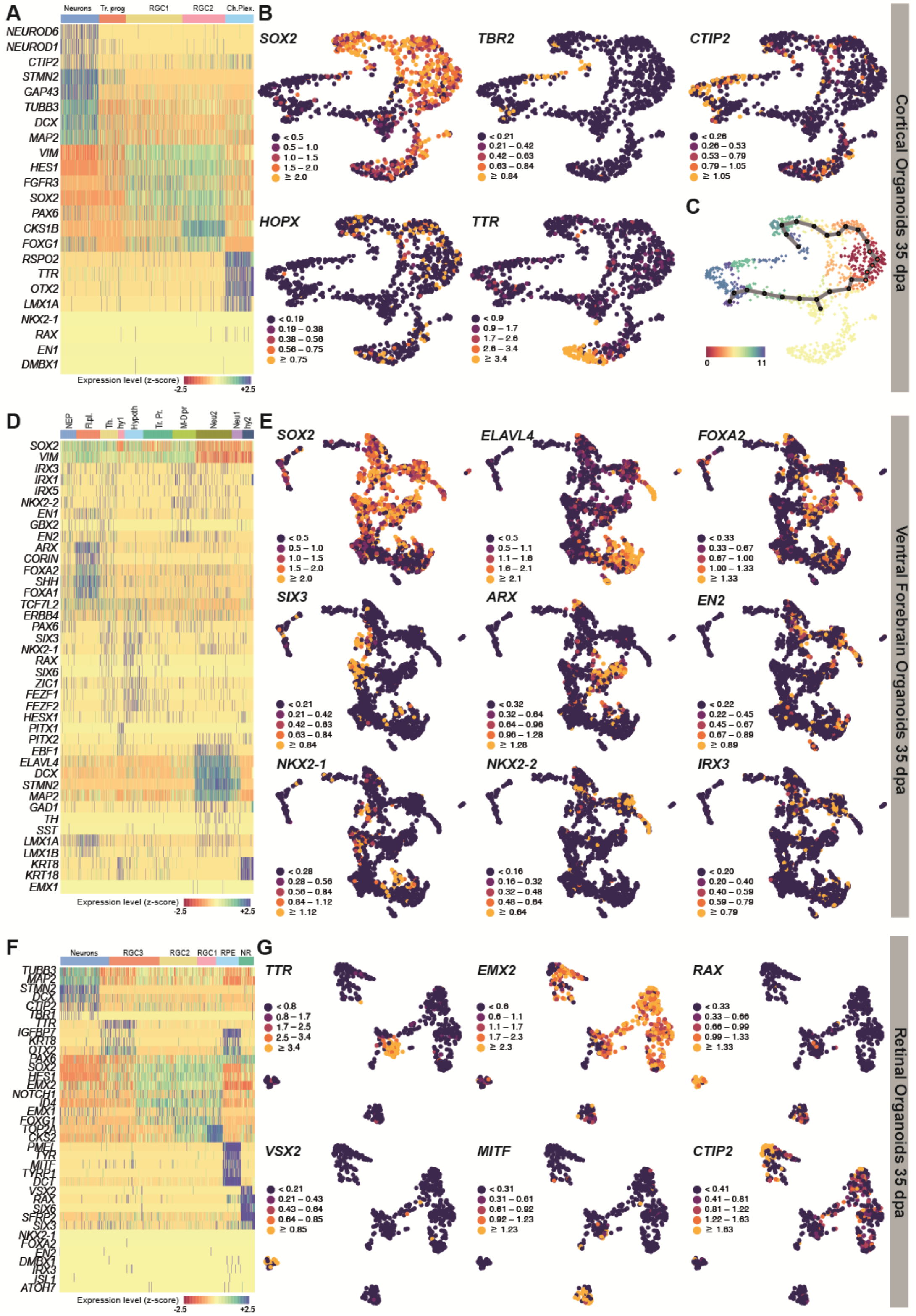


**Supplementary Figure S5. Single cell RNAseq characterization of human COs, VFOs and ROs. A, D,F.** Heatmaps display log-normalized expression values of regional or cell identity marker genes for each Leiden clusters of the COs, VFOs or ROs datasets. The color scale represents capped ($95^{th}$ percentile) Z-score values for each gene. **B,E,G.** Continuous embedding UMAP plots displaying display the expression of selected gene across all cells of two-dimensional UMAP embedding (Fig. 2E). The color scale is set independently for each gene, with high values capped at the 95th percentile to minimize the influence of outliers.

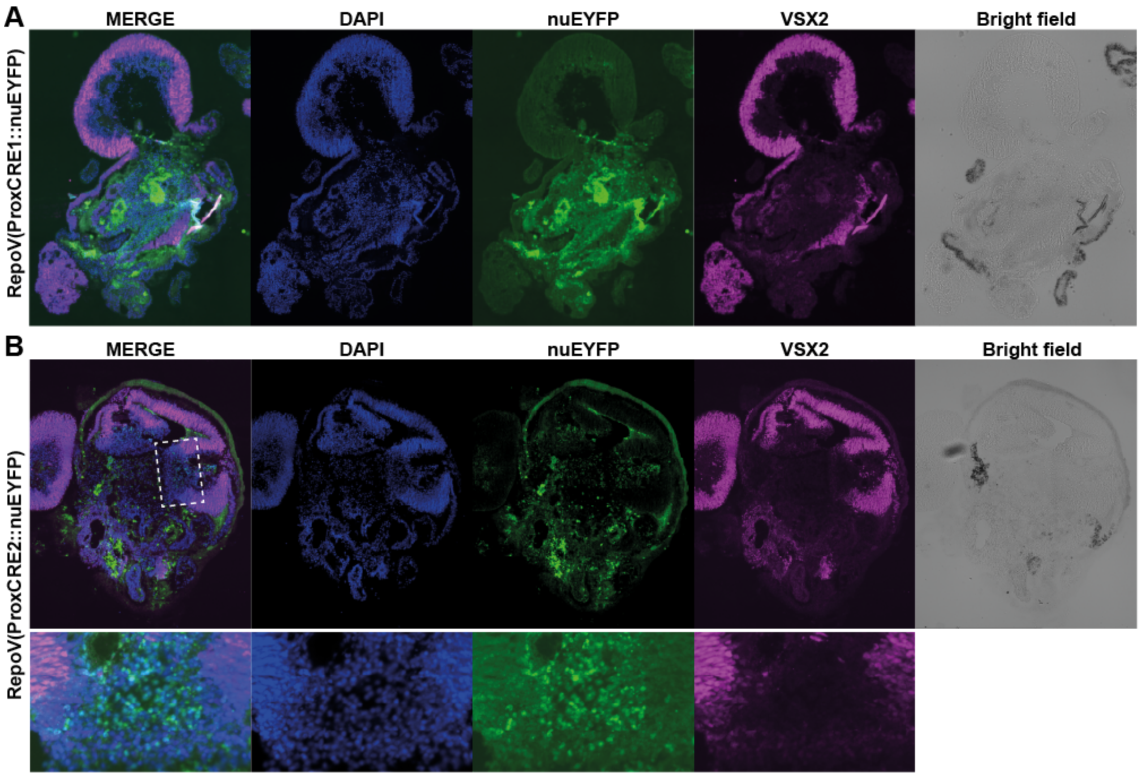


**Supplementary Figure S6.** ***FOXG1*** **ProxCRE1 and ProxCRE2 transgene activity in human ROs. A-B**. Fluorescence and bright field microscopy images of cryostat sections from transgenic ROs showing nuEYFP reporter expression and stained for the neuroretina marker VSX2. Of note ROs are characterized by unspecific green fluorescent signal in their inner portion, likely caused by extracellular matrix components.
