## Supplementary Information for "ORGANOID-BASED ASSESSMENT OF ENHANCER ACTIVITY VIA RECOMBINATION-MEDIATED TRANGENESIS"

### SUPPLEMENTARY RESULTS

#### Characterization of COs, VFOs and ROs.

In all protocols, iPSCs were first aggregated into undifferentiated embryoid bodies (EBs), followed by neural induction (**Fig. 2A**). COs were specified toward dorsal telencephalic identity by inhibiting BMP, TGF-β and WNT signaling, whereas VFOs were induced by BMP and TGF-β inhibition combined with SHH activation. In both cases, subsequent Matrigel embedding promoted the formation of evaginating neuroepithelial vesicles (black arrowheads in **Fig. 2B**), reflecting neuroectodermal expansion. In contrast, ROs underwent spontaneous neural induction, intrinsic self-organization and regional patterning of anterior neuroectoderm, through sequential suspension and adherent culture steps, rather than by morphogen-guided specification (**Fig. 2A,** (Cowan et al. 2020)). This process ultimately leads to the formation of refractile neural-retina like structures (white arrowheads in **Fig. 2B**) flanked by patches of retinal pigmented epithelium (red arrowheads) and emerging from opaque, apparently unstructured aggregates (asterisks).

For COs, RT-qPCR analysis (**Fig. 2C**) revealed strong *PAX6* upregulation compared with EBs, whereas *NKX2-1* and *RAX* showed only modest increases, consistent with dorsal telencephalic specification. Conversely, *NKX2-1* was strongly upregulated in VFOs, with only low enrichment of *PAX6*. *FOXG1*, expressed in both dorsal and ventral telencephalon, was strongly induced in both organoid types. Instead, ROs exhibited robust *RAX* upregulation, contrasting with its limited enrichment in COs and VFOs (**Fig. 2C**). They also expressed high levels of *PAX6* and *FOXG1*, but not of *NKX2-1*, consistent with the expression of these genes in vivo. Immunofluorescence analysis confirmed that CO neurogenic structures comprise PAX6+ radial glial cells forming an in vivo-like ventricular zone, a subventricular zone containing TBR2+ intermediate progenitor cells, and a cortical plate-like structure populated by TBR1+, CTIP2+ and SATB2+ cells (**Fig. 2D** and **Supplementary Fig. S3A,B**). These findings support the generation of both deep- and upper-layer glutamatergic neurons in our COs. Similarly, immunofluorescence analysis confirmed the presence of FOXG1+ and NKX2.1+ domains in our VFOs, together with strong ASCL1 and GAD67 expression, consistent with the specification of ventral forebrain identities and the generation of GABAergic inhibitory neurons (**Fig. 2D** and **Supplementary Fig. S3B**). Finally, neuroretina-like structures in ROs displayed strong PAX6, RAX and VSX2 staining (**Fig. 2D** and **Supplementary Fig. S4**). Retinal pigment epithelium (OTX2+) and optic stalk (PAX2+) domains were also identified in our ROs (**Supplementary Fig. S4**). Notably, we observed patches of PAX6+ RAX− progenitor cells, as well as TBR1+ cells, which tended to localize within the internal, less organized regions of the organoids (**Supplementary Fig. S4**). These observations suggest the presence of cells with cortical identity in our RO cultures, although they do not appear to organize into well-defined neurogenic structures comparable to those observed in COs. At later stages (56dpa) we observed the emergence of retinogenesis markers, as exemplified by the detection of BRN3B + Retinal Ganglion cells towards the lumen of the neurogenic structures and of the CRX+ photoreceptors at the outside (**Supplementary Fig. S4E**).

Single-cell RNA sequencing largely confirmed the regional identity of COs, VFOs and ROs (**Fig. 2E** and **Supplementary Fig. S5**). COs displayed transcriptional signatures characteristic of cortical radial glial cells (*SOX2*, *PAX6*, *NES*) and differentiating glutamatergic neurons. Most neuronal cells corresponded to deep-layer populations (*DCX*, *CTIP2*, *NEUROD4/6*), reflecting the still incipient upper-layer neurogenesis phase at the organoid stage analyzed (35 days post-aggregation), approximately corresponding to the E12.5 mouse cortex. Although intermediate progenitors and basal radial glial cells were not resolved as distinct clusters, we detected *TBR2*+ and *HOPX*+ cells, respectively, suggesting that COs recapitulate the diversity of neural progenitor populations found *in vivo*. In fact, trajectory analysis suggests two developmental routes: direct neuronal differentiation from *SOX2*+ radial glia and indirect neurogenesis involving *TBR2*+ cells (**Supplementary Fig. S5C**). We also identified a choroid plexus cluster (*TTR*, *OTX2*, *LMX1A+)* and little or no expression of ventral forebrain (*NKX2-1*, *RAX*) or caudal (*EN1*, *DMBX1*) markers (**Supplementary Fig. S5A,B**).

In contrast, VFOs displayed a broader diversity of clusters, including *SOX2*+/*VIM*+ progenitors with signatures of different ventral forebrain domains (**Fig. 2E** and **Supplementary Fig. S5D,E**), while dorsal telencephalic markers were largely absent. These populations included *FOXA1/2*+, *CORIN*+ and SHH+ floor plate cells, as well as *NKX2-1*+/*SIX3/6*+/*RAX*+ hypothalamic progenitors, consistent with the SHH activation used in this protocol. Additional progenitor clusters displayed diencephalic identities, although heterogeneous marker expression suggests that finer subdivisions could not be resolved at the sequencing depth and cell numbers analyzed. For example, a meso-diencephalic progenitor cluster contained partially distinct *NKX2.2*+, *IRX3*+ and *EN1/2*+ populations, potentially reflecting prethalamic, thalamic and mesencephalic identities. Although an MGE-like precursor cluster was not clearly resolved, qPCR and immunofluorescence detection of *FOXG1* and *NKX2-1*, together with neuronal markers, support the presence of this subpallial domain (**Fig. 2C,D** and **Supplementary Fig. S3C,D**), as previously reported (Bagley et al. 2017). We also identified two main clusters of differentiating ventral forebrain neurons, expressing markers including *GAD*, *SST* and *TH*. As in COs, the relatively low proportion of terminally differentiated neurons is consistent with the early neurogenic stage analyzed (**Fig. 2D** and **Supplementary Fig.**

**S5D**). Finally, we detected SOX2− epithelial-like precursors expressing *KRT8/18*, potentially corresponding to hypophyseal derivatives, as suggested by *PITX1*/*2* expression (**Supplementary Fig. S4D**). Although a detailed characterization of VFO populations was beyond the scope of this study, these results support the representation of ventral telencephalic, hypothalamic and diencephalic progenitors and their derivatives, with possible contributions from ventral mesencephalic domains.

Finally, scRNA-seq analysis of ROs confirmed the presence of neural retinal cells (*SOX2*, *PAX6*, *VSX2*, *RAX*, *SIX6*) and retinal pigment epithelium (*OTX2*, *MITF*, *PMEL*, *TYR*) (**Fig. 2D** and **Supplementary Fig. S5F,G**), consistent with our immunofluorescence results. Interestingly, we also detected clusters expressing cortical progenitor markers (*EMX1/2*, *PAX6, SOX2*, *VIM*), differentiating cortical neuronal markers (*MAP2*, *STMN2*, *DCX*, *CTIP2*, *TBR1*) and choroid plexus markers (*TTR*, *OTX2*, *LMX1A*) (**Supplementary Fig. S5F,G**). No other prosencephalic identities were clearly detected, as indicated by the absence of markers such as *NKX2-1* and *GSX2*. Retinal neurons expressing *ATOH7*, *ISL1* and *POU4F2* were also absent at this stage, consistent with our immunofluorescence results. Although cortical identity was not reported in the original RO study (Cowan et al. 2020), other studies describe the concomitant specification of eye-field and telencephalic identities under similar neural induction conditions (Gabriel et al. 2021; Liu et al. 2023).

### SUPPLEMETARY MATERIAL AND METHODS

#### Generation of cortical and ventral organoids

Human induced pluripotent stem cells (hiPSCs) were dissociated into individual cells using Gentle Cell Dissociation Reagent (STEMCELL Technologies) following a 8-10 min incubation at 37°C. The cell suspension was transferred to a conical tube containing eTeSR with 10mM Y-27632 and centrifugated 5 minutes at 200xg. Cells were resuspended in 1ml of aggregation medium (AM) consisting of eTeSR supplemented with 20 μM Y-27632 and counted using Trypan Blue to exclude the dead cells. Only preparations exhibiting >90% viability were used for organoid generation.

*Embryoid body formation* For embryoid body (EB) formation, approximately 7,000 viable cells were seeded in 80 μL of AM per well in ultra-low attachment 96-well U-bottom plates. Plates were centrifuged at 200×g for 2 min and incubated at 37°C in a humidified 5% $CO_2$ atmosphere. After 24 h, 80 μL of fresh eTeSR medium was added to each well. EBs were maintained in aggregation medium until reaching a diameter of approximately 400-600 μm, typically by day 3 post-aggregation. At this point we started the neural induction.

*Neural induction* EBs were cultured in neural induction medium (IM) composed of DMEM/F12 supplemented with N2 (1×), NEAA (1×), GlutaMAX (1×), heparin (1 μg/mL), LDN193189 (0.2 μM), and A83-01 (0.5 μM). Prior to transfer, EBs were washed once with DMEM/F12 to remove residual eTeSR medium. Subsequently, one or two EBs were placed in each well of anti-adherent 24-well plates containing 0.8 mL of IM and maintained at 37°C under gentle orbital agitation. To promote *dorsal/cortical specification*, IWR-1-endo was included in the IM at a final concentration of 3 μM from the beginning of neural induction. In contrast, *ventral patterning* was induced by the addition of the SHH pathway agonist SAG (0.1 μM) 24 h after the initiation of neural induction. In our experience, by day 3 after neural induction, organoids consistently developed a neuroepithelial structure that could be readily identified as an optically translucent rim surrounding the EB. Upon the appearance of this characteristic morphology, organoids were transitioned to the neural expansion phase.

*Organoid embedding and expansion* Neural induced organoids were embedded in Geltrex (Thermo Fisher Scientific). Individual EBs were transferred onto sterile parafilm molds and embedded in 20 µL Geltrex droplets. After polymerization at 37°C for 20-25 min, droplets were carefully transferred into 6-well plates containing 3 ml of expansion medium (EM). EM consisted of a 1:1 mixture of DMEM/F12 and Neurobasal medium supplemented with N2 (0.5×), B27 minus vitamin A (1×), GlutaMAX (1×), NEAA (0.5×), β-mercaptoethanol (192.5 µM), and insulin (2.5 µM). Approximately 5-8 organoids were maintained per well under orbital agitation at 37°C, and medium was replaced every 3 days. Organoids were cultured in EM for 3-4 days until the appearance of prominent neuroepithelial buds.

*Organoid maturation.* At this point, organoids were transferred to maturation medium (MM), consisting of a 1:1 mixture of DMEM/F12 and Neurobasal supplemented with N2 (0.5×), B27 containing vitamin A (1×), GlutaMAX (1×), NEAA (0.5×), β-mercaptoethanol (192.5 µM), and insulin (2.5 µM). Cultures were maintained on an orbital shaker at 37°C and 5% $CO_2$, with complete medium changes performed every 3 days until day 20 post induction when the medium changes were performed every 2 days. From 35 dpa, maturation medium was additionally supplemented with 20 ng/mL brain-derived neurotrophic factor (BDNF) and 20 ng/mL glial cell line-derived neurotrophic factor (GDNF). Organoids were maintained under these conditions until the desired experimental endpoint

**KEY RESOURCES TABLE**

| REAGENT or RESOURCE | SOURCE | IDENTIFIER / CAT. No. |
|---|---|---|
| eTeSR | StemCell | 100-1215 |
| DMEM F12 | Gibco | 11320033 |
| NeuroBasal | Gibco | 10888022 |

| | | |
|---|---|---|
| Glutamax | Gibco | 35050038 |
| NEAA | Gibco | 11140050 |
| 2-mercaptoethanol | Gibco | 21985023 |
| Y-27632 (Dihydrochloride) 5mg | StemCell | 72304 |
| LDN193189 (2HCl), 10 mg | StemCell | A72147 |
| IWR-1-endo, 5 mg | StemCell | 72562 |
| 83-01, 5 mg | StemCell | 72022 |
| SAG | StemCell | 73412 |
| Heparin 0,2% | StemCell | 07980 |
| Insulin solution human sterile-filtered | Sigma | I9278-5ML |
| Geltrex™ Flex LDEV-Free hESC-Qualified | Gibco | A4000046803 |
| Microplate 96 well, PS, U-BOTTOMCell repelent Surface | Greiner | 650970 |
| Anti-adherence rinsing solution | StemCell | 07010 |
| Gentle cell dissociation reagent | StemCell | 100-0485 |
| N2 | Gibco | 17502048 |
| B27 | Gibco | 17504044 |
| B27 without vitamin A | Gibco | 12587010 |
| hBDNF | StemCell | 78005 |
| hGDNF | StemCell | 78058 |
